# Multiparametric Assessment of TNNI3 Variant Phenotypes in Human iPSC-Cardiomyocytes Correlates with Disease Severity in Patients

**DOI:** 10.64898/2025.12.13.691658

**Authors:** David W Staudt, Peter PQ Tran, Brendan J Floyd, Kyla Dunn, Dongju Han, Xiomara Carhuamaca, Ricardo Serrano, Anna P Hnatiuk, Seyun Bang, Victoria N. Parikh, Euan A. Ashley, Mark Mercola

## Abstract

**Background:** The routine genetic testing of cardiomyopathy patients has significantly accelerated the identification of causative cardiomyopathy variants. However, translating these genetic insights into effective patient management poses significant challenges, since the impact of gene variants on physiological function and clinical outcomes is not yet fully understood. Therefore, there is an urgent need for large-scale methods to assess the effects of genetic variants on cardiomyocyte physiology and to establish correlations between functional phenotypes and clinical severity.

**Methods:** We developed a high throughput imaging platform to measure force generation and calcium handling throughout the cardiac cycle of human induced pluripotent stem cell-derived cardiomyocytes (hiPSC-CMs). By expressing variants of a sarcomeric protein [cardiac Troponin-I (TNNI3)] in a healthy genetic background, we were able to assess sarcomeric calcium sensitivity as well as systolic and diastolic function. Analysis of these parameters distinguished subgroups of variants, and permitted the correlation of *in vitro* physiological effects with a measure of disease severity in a single-center cardiomyopathy cohort.

**Results:** Combining contractile force and calcium cycling measurements accurately distinguished known pathogenic from non-pathogenic *TNNI3* variants and also revealed pathogenicity of two variants of unknown significance (VUS) that occurred in two families, suggesting the ability to prospectively discern pathogenicity. Clustering of *TNNI3* variants based on quantitative physiological phenotypes identified subgroups that correlated with age of disease onset across a well-characterized cardiomyopathy patient cohort, showing clinical relevance of the *in vitro* phenotypes. Interestingly, normalized measures of *in vitro* diastolic function correlated with age of onset (R^2^ = 0.6), but calcium sensitivity, which accurately predicted pathogenicity, did not translate into disease severity.

**Conclusions:** A high throughput *in vitro* platform that measures multidimensional cardiomyocyte function can link subgroups of human genetic variants in TNNI3 with differential patient outcomes. Comprehensive determination of variant effects on disease-relevant cardiomyocyte function will help classify variants into different pathogenic mechanisms leading to variable disease severity, and potentially lead to class-targeted ameliorative strategies.

## Introduction

Inherited cardiomyopathies cause significant morbidity and mortality secondary to impairment of cardiac function^1–5^. Genetic testing of cardiomyopathy patients has entered routine clinical practice to refine diagnostic precision and aid in personalizing therapies^6,7^. However, current clinical decision making is based on relatively few known pathogenic gene variants, and even for these, there is little information regarding the impact on cardiomyocyte function on clinical severity. Moreover, most gene variants detected in patients are of uncertain significance (VUS) because of insufficient clinical and experimental data associating them with disease^8–10^. Consequently, genetic test results are often inconclusive; and when likely pathogenic or pathogenic variants are identified, they provide positive molecular diagnoses but little prognostic insight to inform patient management.

Human induced pluripotent stem cell-derived cardiomyocytes (hiPSC-CMs) carrying cardiomyopathic protein variants can recapitulate contractile phenotypes consistent with clinical phenotypes, and therefore offer a means to correlate genetic variants with clinical outcomes^11–15^. However, prior studies focused either on only a few variants at a time, or on univariate surrogate markers of muscle function, thus providing too little information to relate the spectrum of genetic variation to disease severity. Here, we developed a platform for high throughput, high-content assessment of individual variants’ effects on systolic and diastolic force production as well as calcium handling in engineered hiPSC-CMs. To investigate the relationship between *in vitro* dysfunction and clinical presentation, we analyzed the multidimensional parameters of *in vitro* physiological function and identified features correlating with clinical findings from a cardiomyopathy patient cohort.

Cardiomyopathies are characterized by their functional phenotype and fall into several distinct categories. Dilated Cardiomyopathy (DCM) is characterized by dilated heart ventricles with poor contractile (systolic) function while Hypertrophic Cardiomyopathy (HCM) presents with thick ventricular walls, normal to increased contractile function and alterations in relaxation and filling (impaired diastolic function). In contrast, Restrictive Cardiomyopathy (RCM) patients have normally sized ventricles but severely impaired diastolic function. Each of these subtypes can be caused by alterations of the cardiac sarcomere. *TNNI3* encodes cardiac troponin I, a central element of the sarcomere that interacts with tropomyosin to inhibit the association of actin and myosin at resting (diastolic) calcium levels^5, 16–20^. Variants in *TNNI3* comprise the most prevalent known genetic lesions associated with RCM but also cause HCM and, less prevalently, DCM^5,16^. At the time of writing, 374 *TNNI3* missense variants have been identified in cardiomyopathy patients and listed in ClinVar^21^. Only 62 are classified as either Pathogenic, Likely Pathogenic, Likely Benign, or Benign, and the majority are VUS. Further, clinically important questions such as why some variants cause HCM, RCM, or DCM, or why some patients present earlier than others, are not fully understood.

Using our platform, we quantified the systolic, diastolic, and calcium cycling effects of a library of *TNNI3* variants in ClinVar on hiPSC-CM function. Derangement of calcium sensitivity and force generation *in vitro* accurately classified variants as HCM/RCM, or DCM-causing, and provided evidence for pathogenicity for VUS seen in two cardiomyopathy families. Multiparametric clustering divided variants into multiple functional clusters, and identified groups of variants that presented at earlier ages of onset in a patient cohort. Lastly, we identified key diastolic metrics that correlate with the observed subgroup differences in severity. Thus, this platform provides a comprehensive tool for assessing the differential effects of cardiomyopathy variants on disease-relevant cardiomyocyte function at scale, and lays the groundwork for identifying functionally and clinically distinct subgroups of cardiomyopathy.

## Methods

### Human iPSC culture and differentiation

Human iPSCs were described previously^22^ and had been generated with approval of the Stanford Institutional Review Board and Stem Cell Research Oversight Committee. Human iPSCs were cultured with daily media changes in E8 medium (Thermofisher) on Geltrex (Thermofisher)-coated 6-well tissue culture plates and incubated at 37 °C and 5% CO2. Cells were passaged with PBS/EDTA and Rock inhibitor Y-27632 (Tocris) every 2-3 days at ∼80% confluency.

hiPSCs were differentiated to cardiomyocytes using previously published protocols^22^. At 90-95% cell confluency, hiPSCs were differentiated into cardiomyocytes following a chemically defined differentiation protocol. In brief, hiPSCs were treated with CHIR-99021 (Tocris) in RPMI 1640 with B27-insulin supplement (Thermofisher) at differentiation day 0. At day 1 and 2, CHIR-99021 was gradually diluted with RPMI 1640/B27-insulin. On day 3, the differentiating cells were treated with Wnt-C59 (Tocris) in RPMI 1640/B27-insulin. On day 5 and 7, full media changes with RPMI 1640/B27-insulin were done. On day 9, media was exchanged with RPMI 1640 with complete B27 supplement (Thermofisher). On day 11, a full media change with RPMI 1640-glucose/B27 was performed for metabolic selection to improve cardiomyocyte purity. On day 14, media was replaced with RPMI 1640/B27 to let cardiomyocytes recover from metabolic selection. On day 15, cardiomyocyte purity was assessed under light microscope and pure cardiomyocyte cultures were dissociated with TrypLE 10X (Thermofisher) and replated onto Geltrex-coated 6-well plates at a seeding density of 3 x 10^6^ cells per well in RPMI 1640/B27 with 10% Knockout Serum Replacement (KOSR, Thermofisher) and Rock inhibitor Y-27632. On day 16, media was changed to RPMI 1640/B27. On day 17, cardiomyocytes went through a second round of metabolic selection with RPMI 1640 minus glucose/B27 and, on day 20, media was changed to RPMI 1640/B27. Cardiomyocytes were afterwards maintained in RPMI 1640/B27 with media changes twice a week.

### Genome editing of human iPSCs

CRISPR-Cas9 knockout of TNNI1 and TNNI3 were done sequentially in a low passage wildtype human iPSC line (15S1).

hiPSCs were plated with E8 media and Rock inhibitor at low density onto Geltrex-coated 6-well plates a day before transfection. On the day of transfection, 20 pmol Cas9-enzyme and 60 pmol sgRNA per 250,000 cells were mixed into final volume of 4 µL and incubated at room temperature. Target cells were dissociated into single cells and resuspended at a concentration of 250,000 cells per 10 µL in Buffer R. 10 µL of target cell suspension was added to 4 µL of Cas9 and sgRNA mix. Target cells were then electroporated in Buffer E with 2 x 1200 V for 20 ms using Neon™ Transfection System (Thermofisher). Electroporated cells were replated into a well of a Geltrex-coated 12-well plate with E8 media and Rock inhibitor. At first passage post-transfection, a few cells were resuspended in 20 µL media and genomic DNA was isolated to assess editing efficiency by PCR amplification of target genomic edit site. PCR products were Sanger sequenced with results analyzed by ICE analysis tool (EditCo). With sufficient editing efficiency confirmed, individual hiPSC clones were isolated and expanded. Genomic DNA was isolated from clones to confirm an edited genotype by PCR amplification of target genomic edit site and Sanger sequencing. Clones were karyotyped for chromosomal abnormalities with KaryoStat assay (ThermoFisher) (Figure S1). Clones with confirmed edited genotypes were further investigated for potential off-target edits predicted by COSMID (https://crispr.bme.gatech.edu/) ^23^. Notably, initial studies shown in Figure 1 were performed in a clone that was subsequently found to contain a small heterozygous deletion in DLGAP2, a neural gene not detected in hiPSC-CMs (Figure S2b, clone 22). A second clone was isolated that did not have this off-target edit (Figure S2c, clone 10). All subsequent experiments in Figures 2-5 were performed with this clone.

**Figure 1:**
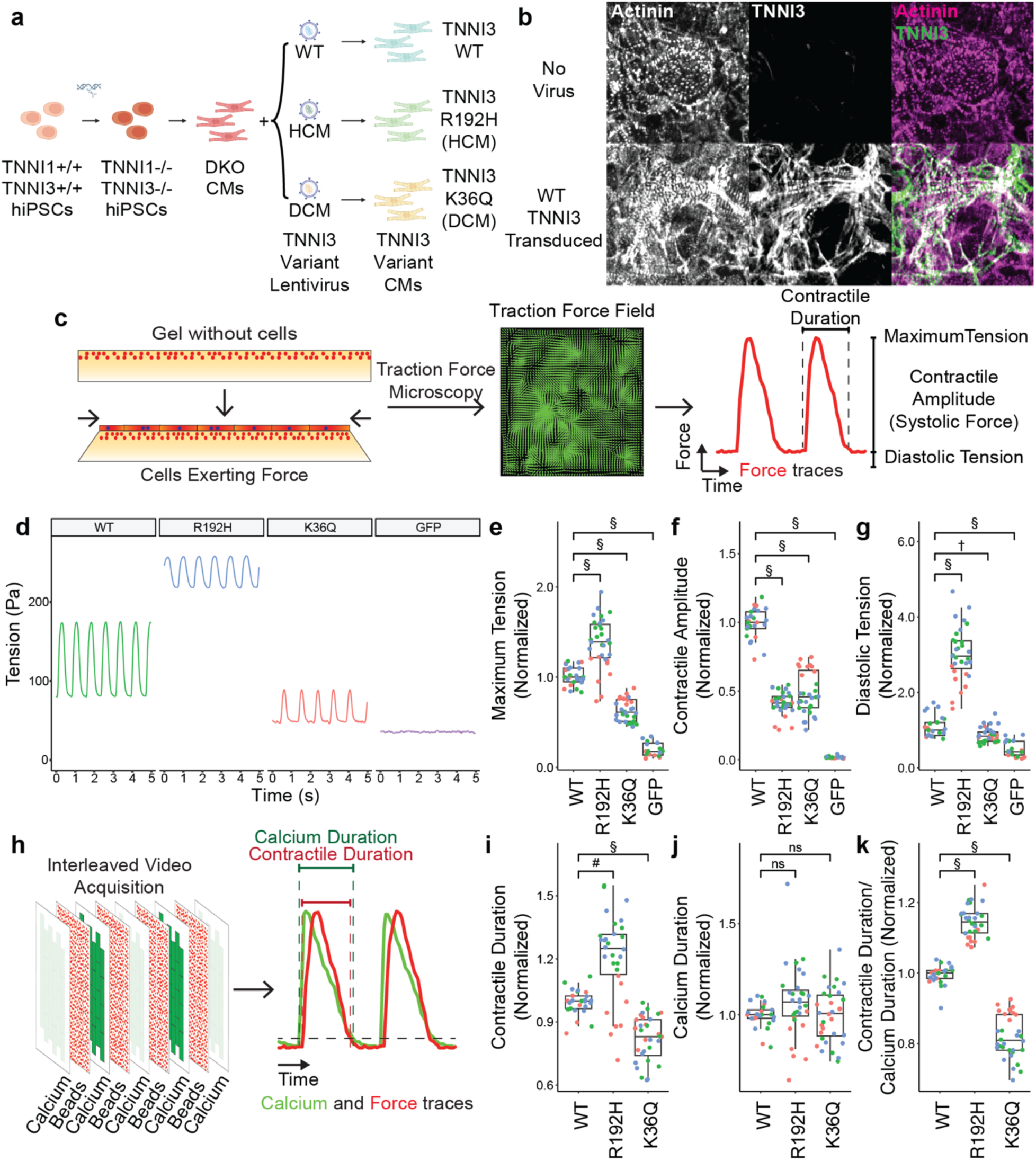
Lentiviral Platform using hiPSC-CMs for Measuring Functional Effects of TNNI3 Mutations. **a.** Schematic of platform showing TNNI1/TNNI3 double knockout cells and lentiviral gene replacement. **b.** Immunostaining reveals disorganized actinin and no TNNI3 expression in untransduced cells, with rescue of sarcomeric structure and TNNI3 function upon viral induction **c.** Schematic of platform for measuring traction forces **d.** Example traces from cells transduced with WT TNNI3, an HCM/RCM mutant (R192H), a DCM mutant (K36Q) or GFP control. **e.** Maximum generated tension, **f.** Systolic contractility, and **g.** Diastolic tension measurements for these variants. **h.** Schematic of interleaved, synchronized calcium and contractility imaging. **i.** Contractile duration, **j.** Calcium duration, and **k.** Contractility/Calcium ratio measurements for these variants. After removing wells without adequate gels, 25 WT wells, 31 R192H wells, 32 K36Q wells, and 18 GFP wells were analyzed across 3 independent differentiations for all variables. Significance calculated using Wilcoxon rank-sum test with holm correction for multiple testing. *p<0.05, †p<0.01, ‡p<0.001, §p<0.0001

**Figure 2:**
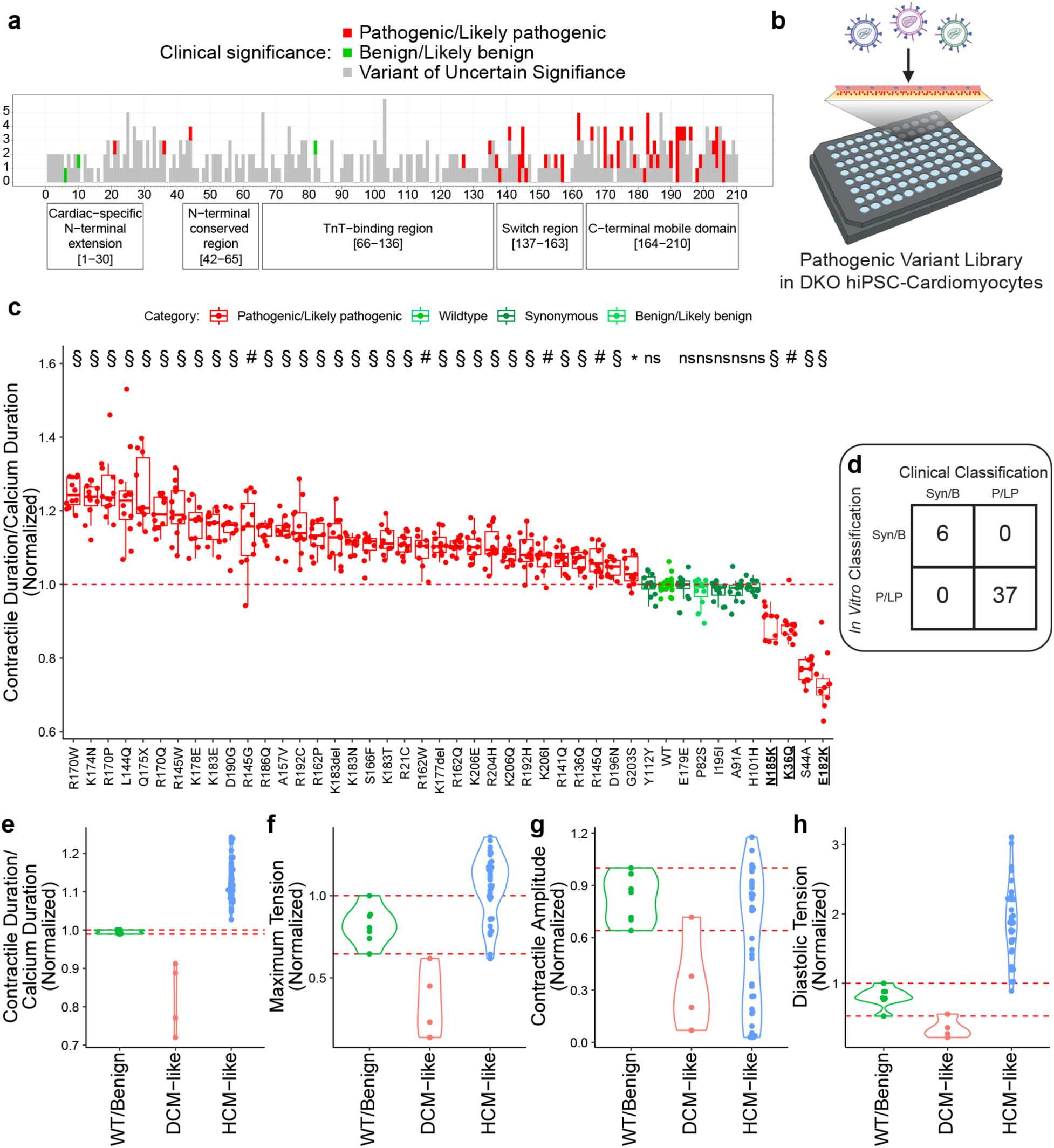
Multidimensional Functional Measurement of a Variant Library. **a.** Histogram showing locations of variants found in ClinVar. **b.** Schematic of Variant library in 96 well plates. **c.** Contractility Duration/Calcium Duration ratio of a 43-member variant library normalized to WT. Pathogenic variants are shown in red, WT, synonymous and benign variants are shown in green. **d.** Comparison of *in vitro* classification of synonymous or benign (Syn/B) variants or pathogenic/likely pathogenic (P/LP) vs annotation in ClinVar **e.** Comparison of WT/Benign, HCM-like, and DCM-like variant Contractility/Calcium Ratio, **f.** Maximum Tension, **g.** Contractile Amplitude, and **h.** Diastolic Tension. Red dotted lines indicate maximum and minimum values seen in WT/Benign variants. A total of 12 replicates/variant and 36 WT replicates from 3 independent differentiations were analyzed across all variables shown. Significance calculated using Wilcoxon rank-sum test with holm correction for multiple testing. *p<0.05, †p<0.01, #p<0.001, §p<0.0001

### TNNI3 variant plasmid cloning

A custom lentivirus transfer plasmid vector that included a CAG promoter, IRES, mCherry-P2A-Neomycin and certain restriction sites was cloned by Gibson assembly. The custom lentivirus transfer plasmid vector was named PTpLV5.

For TNNI3 variant cloning, PTpLV5 was first digested with EcoRI and BamHI (NEB). WT TNNI3 was PCR-amplified from a cDNA template and cloned into the EcoRI/BamHI cut site of PTpLV5 by Gibson assembly (NEBuilder® HiFi DNA Assembly Master Mix, NEB). Pathogenic, Likely Pathogenic, and benign variants were identified from ClinVar. The five most common synonymous variants identified in the gnomAD database were also included as controls. Mutant TNNI3 variants were generated by PCR-amplification with Q5 polymerase (NEB) of two DNA fragments with variant-specific mutagenic primers that each was paired with a universal N-terminus or C-terminus primer. Together, two DNA fragments would result in full length mutant TNNI3 DNA that was cloned into the EcoRI/BamHI cut site of PTpLV5 by 3-fragment Gibson assembly. In the case of the mutation being in either the N-terminus or C-terminus of TNNI3, a single variant-specific mutagenic primer was paired with a universal N-terminus or C-terminus primer for PCR amplification of full length mutant TNNI3 DNA that would be cloned into the EcoRI/BamHI cut site of PTpLV5 by Gibson assembly.

The Gibson assembly mixes were transformed into NEB Stable E. Coli and plated onto agar plates with ampicillin for overnight incubation at 30°C. The next day, colonies were picked and grown in LB media with carbenicillin in a shaking incubator at 220 rpm and 30°C overnight. Plasmid DNA was harvested by miniprep. All plasmids generated were sequence verified by whole plasmid sequencing (Plasmidsaurus).

### Lentivirus production and titering

Lenti-X 293T HEK cells (Takara) were cultured in T75 flasks with DMEM/F12 and 10% FBS (Thermofisher). Cells were passaged at ∼80% confluency with TrypLE 1X. At the day of transfection and 80-90% cell confluency, full media change with DMEM/F12 and 10% FBS was done in the morning. In the afternoon of the same day, 1.72 pmol psPAX2 plasmid, 0.95 pmol pMD2.G plasmid, and 2.17 pmol transfer plasmid were mixed with Opti-MEM (Thermofisher) in a microcentrifuge tube to 500 µL final volume. In a separate microcentrifuge tube, 1 mg/mL Polyethylenimine (PEI, Polysciences) was diluted in Opti-MEM to 500 µL final volume so that the ug DNA:µg PEI was 1:3. The PEI mix was added dropwise to the plasmid mix while gently flickering the tube followed by 20-minute incubation at room temperature. After incubation, the combined plasmid and PEI mix was gently added to the media of the target T75 flask of Lenti-X 293T. In the morning of day 1 post-transfection, full media change with DMEM/F12 and 10% FBS was done. On day 4 post-transfection, the media containing lentivirus was harvested and 1 volume of Lenti-X concentrator (Takara) was added to 3 volumes of media. The mixture was incubated at 4 °C between 1 hour to overnight. The cooled harvested mixture was centrifuged at 1,500 x g and 4 °C for a minimum of 45 minutes. After centrifugation, the supernatant was aspirated, and the pellet was resuspended in RPMI 1640/B27 with 10% KOSR and Rock inhibitor in 1/100 of the initial volume. Aliquots of concentrated lentivirus were made and stored at -80 °C.

### Lentiviral Titering

For experiments shown in Figure 1, a virus containing GFP was reverse transduced onto 15,000 DKO-CMs in a serial dilution. On day 4 post replating, cells were stained with Hoechst (Thermofisher) and imaged on the Kinetic Image Cytometer. Titer was determined by calculating the percent of GFP positive cells. GFP and TNNI3 viral titers were then measured using p24 ELISA (Takara), and normalized to the GFP control.

For all subsequent experiments, viruses were designed to express mCherry for direct titering. Concentrated lentivirus stocks were functionally titered on DKO-CMs in 384-well plates. Cells were reverse transduced by replating cells on top of serial diluted lentivirus in RPMI 1640/B27 with 10% KOSR and Rock inhibitor. On day 4 post-replating, cells were stained with Hoechst (Thermofisher) and MitoTracker DeepRed (Thermofisher) for fluorescent imaging. To determine the functional titer, mCherry-positive cells were counted and compared to the number of total Hoechst-positive nuclei. A reference virus was titered on each plate to allow for normalization and calculations of relative titers from plate to plate.

### Functional contractility and calcium assay

On the day before transduction, 96-well plates with 8 kPa polyacrylamide hydrogels embedded with blue, fluorescent beads (Matrigen) were coated with Geltrex (Thermofisher). Well-center images of the bead positions were taken with Kinetic Image Cytometer for quality control. Wells with beads that were not in focus were marked and not imaged downstream and a plate map was designed accordingly. Geltrex-coated soft substrate plates were then incubated overnight at 37 °C and 5% CO2.

On the morning of transduction, reference images of the beads at five different well positions were taken for post-analysis. In a 96-well deep-well plate, TNNI3 variant lentiviruses at the same relative titers were mixed with RPMI1640/B27 with 10% KOSR and Rock inhibitor. TNNI3 variant lentiviruses were diluted to reach a final MOI of ∼2.5 to achieve >90% infected cells in each well. Normalized lentiviral mixtures were transferred to polyacrylamide hydrogel-containing 96 well plates according to the designated plate map. After successful transfer of lentiviruses, the plates were briefly spun down at 200 x g for 10 seconds and then returned to the incubator. Day 23 DKO-CMs were dissociated with TrypLE 10X and 150,000 cells per well were replated onto the four plates carrying lentivirus for a reverse transduction. The cells were allowed to settle in the incubator for 10 minutes before being spun down at 200 x g for 10 seconds and then returned to the incubator.

On day 1 post-transduction, RPMI 1640/B27 with final 1% penicillin/streptomycin was added to each well. On day 5, half media change with RPMI 1640/B27 with 1% penicillin/streptomycin was performed.

On day 7 post-transduction, the cells were washed with 3x half media changes of FluoroBrite DMEM and a final 1 x half media change with FluoroBrite DMEM and Cal-520 AM (AAT Bioquest) at a final concentration of 625 nM. The cells were then incubated for 1 hour at 37 °C and 5% CO2. After incubation, the cells were washed with 4 x half media changes of FluoroBrite DMEM and then returned to the incubator for 30 minutes. All media used was preheated in a water bath to 37 °C. After incubation, transduced DKO-CMs were imaged on a Kinetic Image Cytometer (Vala) with interleaved acquisition of the movement of the fluorescent beads (contractility) and fluorescence intensity of Cal-520 (calcium transients). Contractility and calcium measurements were extracted from the raw movies using in house Matlab scripts^24^.

### Multivariable Clustering of Variants

One hundred random forest models classifying pathogenic vs non-pathogenic variants were created incorporating all 34 measured functional variables. Importance scores were normalized for each model and then averaged across models to create an average normalized score. This was used to rank the variables in order of importance. The top 5 variables were chosen based on the observed drop-off in the normalized importance, and these were used for hierarchical clustering using Ward’s method. The optimal number of clusters was determined by identifying the cluster number producing the highest silhouette score.

### Immunostaining

DKO-CMs were washed twice with PBS and fixed with 4% paraformaldehyde for 15 minutes. Cells were permeabilized and blocked with 0.3% Triton-X and 5% BSA in PBS for one hour, after which they were stained with the primary antibodies at 1:1000 dilution with 0.3% Triton-X and 1% BSA in PBS overnight. Cells were washed 3 times with PBS and then stained with secondary antibodies (Thermofisher) at a 1:200 dilution for 2 hours. Cells were then washed with PBS and kept until imaging. Primary antibodies used were: polyclonal rabbit-anti-TNNI3 (Abcam ab47003), and mouse monoclonal anti-sarcomeric alpha-actinin (Thermofisher MA1-22863).

### Patient Data

Probands evaluated at Stanford Medical Center or Stanford Medicine Children’s Health who were found to have variants in TNNI3 on routine clinical sequencing were identified retrospectively, as approved by the Stanford Research Compliance Office, IRB protocol 4237. Deidentified clinical data were collected. Variants were classified according to the American College of Genetics and Genomics criteria. With the exception of a shared list of variants, clinical and in vitro data were initially collected and analyzed by separate researchers, and were only combined once all data collection had been completed to avoid experimental bias.

### Data Analysis and Reproducibility

Data were compiled and analyzed via custom R scripts. As non-normal distributions were routinely observed in all variables measured when combining multiple batches, measurements were compared using a Wilcoxon rank-sum test. Statistics were adjusted for multiple testing using a Holm-Bonferroni correction accounting for all variable comparisons shown. Linear regressions, coefficients of determination, and model significance were calculated using R’s built-in functions. To control for batch effects, all experiments were repeated with at least three independent differentiations of hiPSC-CMs.

## Results

### Implementation of a Functional Phenotyping Platform for Evaluating *TNNI3* Variants

We set out to develop a scalable, functional assay platform for parallel screening of multiple *TNNI3* cardiomyopathy variants. First, we created a hiPSC line that lacked any troponin I. To do so, we used CRISPR/Cas9 to knock out both *TNNI3* and *TNNI1* (Figure 1a), as hiPSC-CMs predominantly express this latter isoform^25–27^. The resulting troponin I double knockout cardiomyocytes (DKO-CMs) had normal karyotype (Figure S1). Despite the absence of troponin I, they differentiated into cardiomyocytes as shown by staining positive for cardiac actinin (Figure 1b). However, they did not assemble normal sarcomeres and did not contract. Lentiviral overexpression of wild-type (WT) *TNNI3* rescued sarcomere formation (Figure 1b). Next, we used this system as a high-throughput platform to measure contractility of *TNNI3* variants in parallel. First, 96 well plates containing 8 kPa polyacrylamide gels with embedded fluorescent microbeads were imaged to determine the initial bead positions (“reference frame”). Cells were then virally transduced and plated onto these plates with one virus used per well. The amounts of virus were carefully normalized prior to the experiment (see Methods). Videos of bead motion caused by beating cardiomyocytes were collected at the same positions, and bead displacement relative to the reference was calculated for every frame using GPU-accelerated particle image velocimetry. Traction force microscopy was then used to calculate forces relative to the no-cell reference frame^24^ (Figure 1c). While DKO-CMs transduced to express an inert eGFP showed no dynamic contractility, cells transduced to express wild-type TNNI3 robustly generated systolic and diastolic forces (Figure 1d-g).

Prototypical pathogenic variants that cause RCM or HCM (R192H)^28, 29^ or DCM (K36Q)^30^ exhibited distinct phenotypes consistent with their in vivo effects: R192H increased both maximum and diastolic tensions compared to WT (Figure 1e and 1g) while K36Q decreased these values (Figure 1e). Interestingly, both variants exhibited decreased generated systolic tension (defined as the maximum minus diastolic tensions, Figure 1f).

### Scalable Measurement of Calcium-Contractility Relationships

Given that altered calcium-contractility relationships are a known pathogenic mechanism for various cardiomyopathies^31, 32^, we incorporated calcium cycling measurements into our platform. By staining cells with a calcium-sensitive dye and acquiring movies in an “interleaved” fashion, rapidly toggling excitation wavelengths to alternate between measurement of the calcium signal and the bead position, we obtained synchronized dynamic measurements of cytosolic calcium and force generation (Figure 1h).

Comparing the kinetics of contractility and calcium in the DKO-CMs revealed that the R192H mutation increased while K36Q decreased total contractile duration compared to WT (Figure 1i). However, neither mutation affected the kinetics of calcium cycling, suggesting intrinsic differences in calcium sensitivity are dominant drivers of the contractile phenotypes in these variants (Figure 1i, j)^30, 33^. Accordingly, the ratio of contractile duration to calcium duration yielded a measurement of contractile dependence on calcium that distinguished R192H and K36Q mutations from each other and from WT (Figure 1k). Specifically, the R192H variant increased this ratio, suggesting increased calcium sensitivity, and conversely the K36Q variant decreased this ratio, consistent with decreased calcium sensitivity.

### Multiparameter Functional Measurement of a Pathogenic Library

ClinVar lists 379 variants in TNNI3, of which 62 are listed as pathogenic and 4 are benign or likely benign (Figure 2a). Using our platform, we assessed a library of *TNNI3* variants listed as pathogenic in ClinVar^21^, and compared them with one benign and five synonymous variants frequently identified in the gnomAD dataset^34^ (Figure 2b). Given that alterations in the calcium-contractility relationship differentiated both HCM/RCM variants and DCM variants from WT in our assay, we investigated whether the contractile/calcium duration metric could distinguish pathogenic variants from benign and synonymous variants. Remarkably, all 37 pathogenic variants assayed showed altered contractile/calcium duration ratios, whereas this ratio was unchanged compared to WT in benign and synonymous variants (Figure 2c-e). Rank ordering variants based on this metric discriminated HCM and RCM from DCM causal variants (Figure 2c). DCM variants previously characterized in the literature (K36Q, N185K^30^, and E182K^35^) exhibited a diminished calcium-contractility duration ratio. The sole exception was S44A, a variant that, to our knowledge, has not been functionally interrogated previously, which is associated with RCM in ClinVar but has a clear DCM phenotype in our *in vitro* assays. We next separated the variants into three groups based on their calcium-contractility relationship: “WT/Benign”, “DCM-like”, and “HCM-like” (Figure 2e). We then compared force generation metrics across these categories. Maximum tension was decreased in DCM-like variants and was normal to increased in HCM-like variants (Figure 2f). Contractile amplitude was decreased in DCM-like variants but showed a wide distribution in HCM-like variants (Figure 2g). Finally, diastolic tension was generally decreased in DCM-like and generally increased in HCM-like variants (Figure 2h). Notably, none of these metrics other than the calcium-contractility relationship perfectly distinguished WT/Benign from HCM-like variants.

### Disambiguation of Clinical Variants of Uncertain Significance

Given that most genetic testing results in VUS, tools are needed to assess their clinical significance and improve clinical diagnosis. To assess the ability of our platform to detect pathogenicity in VUS, we selected 9 *TNNI3* VUS probands who had undergone comprehensive cardiac evaluations in the Stanford Inherited Cardiac Disease clinic and tested their *TNNI3* variants in our platform. Of these variants, 7 showed a statistically significant deviation in their calcium-contractility duration ratios compared to wild-type TNNI3 (Figure 3a), and 3 were within the range observed for confirmed pathogenic variants (RR145-146dup, R69S, and Q48P, Figure 3b). We focused on these outlier variants to determine whether our *in vitro* phenotyping could accurately classify each variant according to the clinical presentation. Due to additional confounding pathogenic mutations in separate cardiomyopathy genes in the patient carrying the R69S variant, we focused the resulting cardiomyopathy phenotyping on families carrying either the RR145-146dup or Q48P variants.

**Figure 3:**
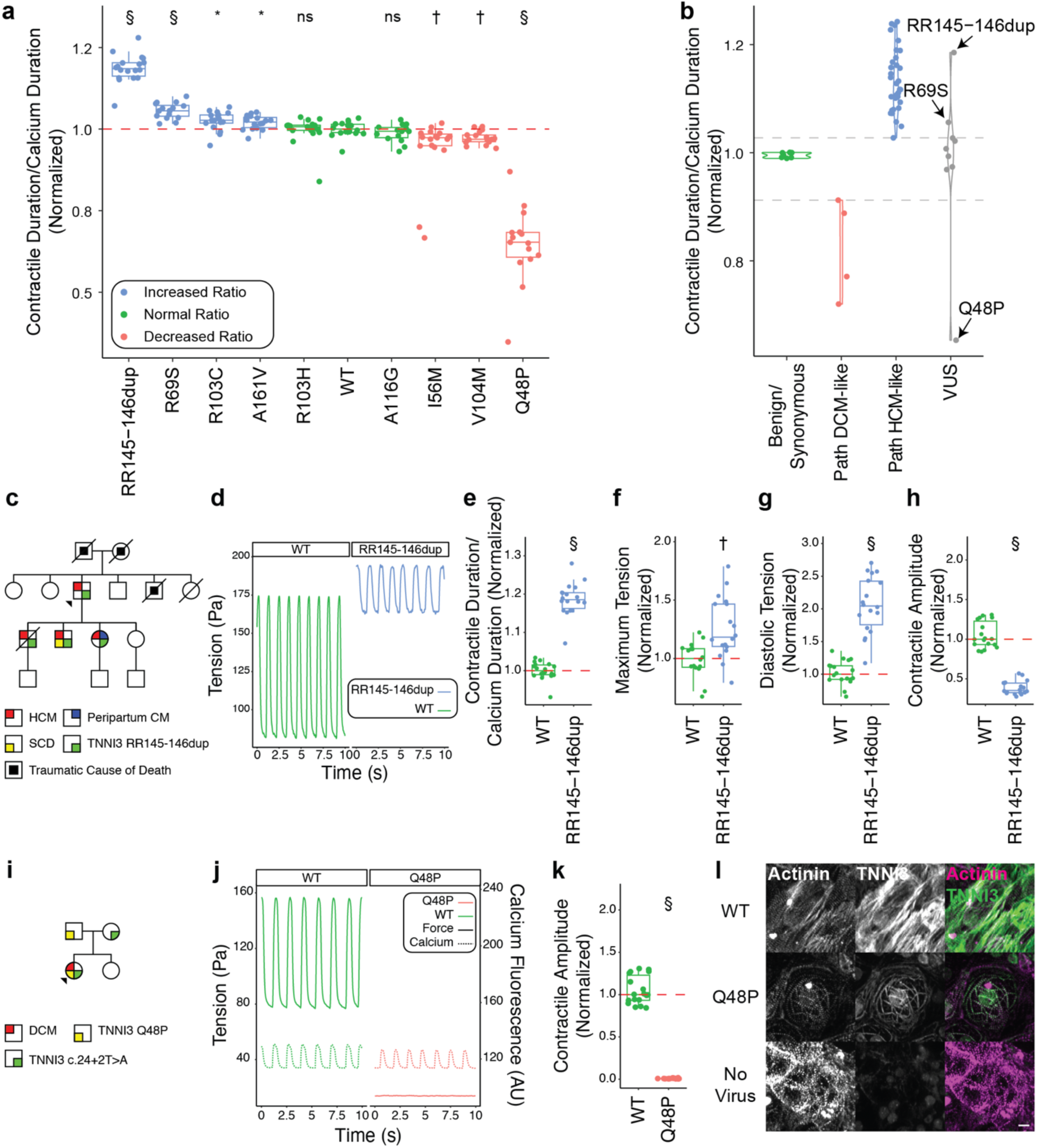
Interrogation of Patient VUS: **a.** Contractility/Calcium ratio of 9 VUS identified in Stanford patients normalized to the median of WT (dashed red line). **b.** Distribution of Contractility/Calcium ratio split by categorization. Specific VUS lying in the pathogenic range are labeled. **c.** Pedigree of Family 1 carrying RR145-146dup variant. **d.** Example traces from WT and RR145-146dup cells. **e.** Contractility/Calcium ratio, **f.** Maximum Tension, **g.** Diastolic Tension, and **h.** Systolic amplitude for the RR145-146dup variant. **i.** Pedigree of Family 2 with Q48P variant. Proband is compound heterozygous for the Q48P and c.24+2T>A variants. **j.** Example traces for Q48P variant. Calcium traces shown with dotted lines. **k.** Systolic amplitude for Q48P variant. **l.** Sarcomeric immunostaining showing disorganized sarcomeres in Q48P mutant. A total of 18 replicates/variant across 3 independent differentiations were analyzed for all variables shown. Significance calculated using Wilcoxon rank-sum test with holm correction for multiple testing. *p<0.05, †p<0.01, #p<0.001, §p<0.0001

Family 1 included four individuals across two generations harboring the RR145-146dup variant, all of whom had HCM (Figure 3c). Functional analysis of this variant revealed a notably increased calcium-contractility duration ratio (Figure 3e), as well as increased maximum and diastolic tensions, and decreased systolic amplitude (Figure 3d-h), consistent with an HCM-like pattern as seen in this family.

In Family 2, the proband presented as an infant with severe DCM (Figure 3i). Clinical genetic testing revealed two variants in *TNNI3*. The first was a known pathogenic splicing variant, c.24+2T>A, inherited from her unaffected mother. This variant has been shown to cause DCM or LVNC in a homozygous state, and led to a complete absence of TNNI3 protein in one homozygous patient^36, 37^. The second variant c.143A>C, p.(Q48P), was classified as a missense VUS. Functional assessment of this second variant revealed a complete absence of force generation, equivalent to that of control DKO-CMs, despite preserved calcium cycling (Figure 3j, k). Immunostaining revealed cells with this variant formed sparse and disorganized sarcomeres compared to WT control (Figure 3l), indicating that this variant is non-functional. Together, these results are consistent with a compound heterozygous loss of TNNI3 function causing DCM in this patient.

### Clustering Analysis to Identify Functional Signatures

The previous analysis focused on rationally chosen metrics based on their interpretability and understandable link to disease phenotypes. However, our platform calculates more than 30 metrics which relate to cardiomyocyte function. Thus, we examined whether other metrics, or combinations thereof, might provide more nuanced information about the myopathic phenotypes of different variants. The first task was to identify which variables would reveal the most information about the observed phenotypes in an unbiased manner. We created a family of random forest classifiers for pathogenicity using all measured and derived variables (34 in total), and extracted the importance metrics for each variable. Averaging these values together created an importance score (Figure 4a). A clear drop-off in this score was observed after the top 5 metrics, so these were chosen for further analysis. The identified metrics included two reflecting calcium sensitivity, both the previously measured contractile duration/calcium duration ratio and the ratio of contractile relaxation time to calcium fall time. Additionally, there were two metrics relating to diastole, namely the ratio of diastolic force to total generated force and the mechanical relaxation rate, and the final metric was the rise time of calcium.

**Figure 4:**
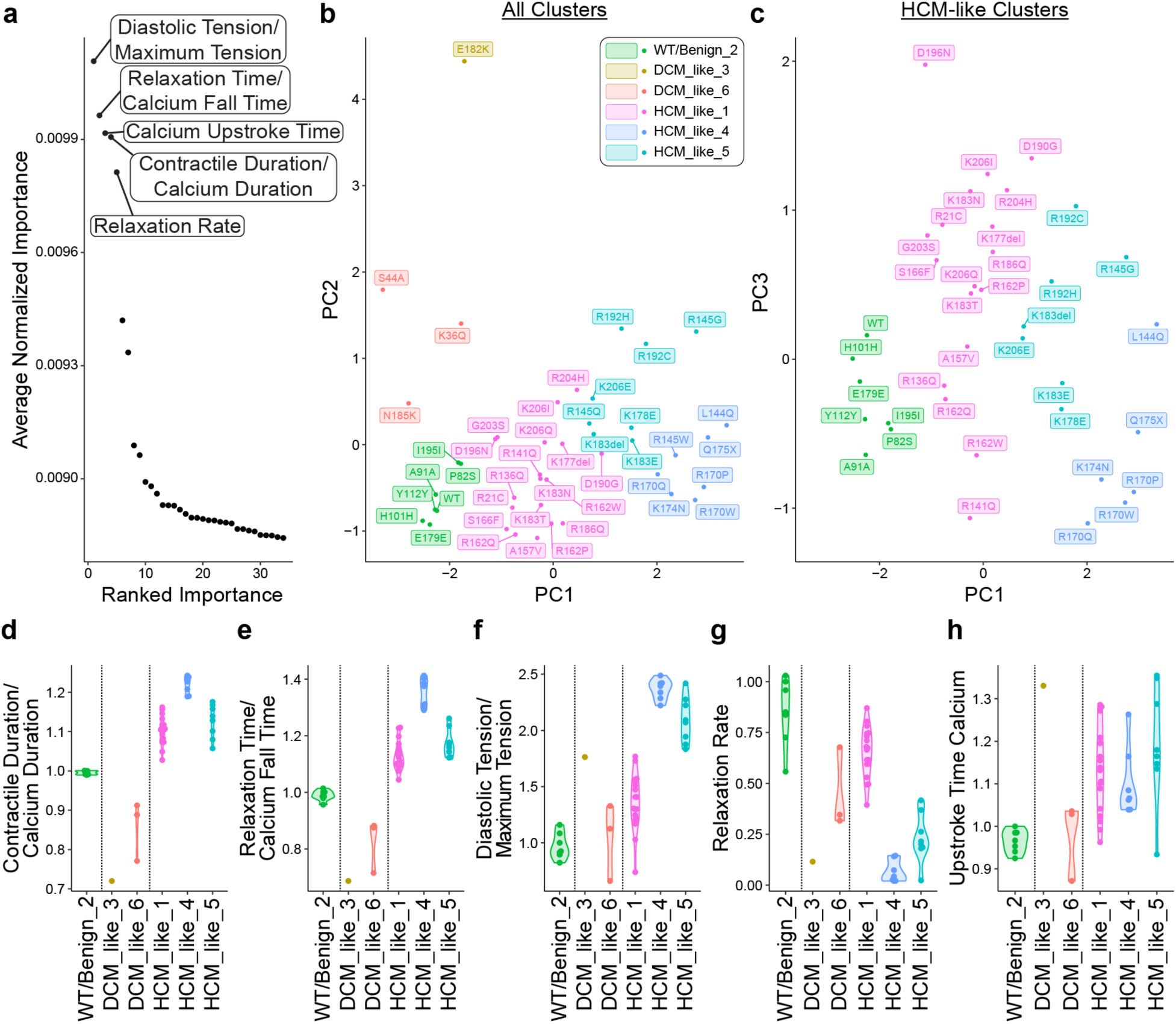
Clustering Analysis of TNNI3 Variants. **a.** Average normalized importance of each variable computed from random forest models. The top five variables chosen for cluster analysis are labeled. **b.** PCA plot of all clusters. **c.** PCA plot showing only WT/Benign and HCM-like clusters. **d.** Comparison of Contractility Duration/Calcium Duration ratio in each cluster. Vertical lines divide WT/Benign, DCM-like, and HCM-like clusters. **e.** Similar comparison of Relaxation Time/Calcium Fall Time, **f.** Diastolic Tension/Maximum Tension, **g.** Relaxation Rate, and **h.** Upstroke Time of Calcium.

We performed hierarchical clustering of variants using these metrics and split the tree into multiple clusters that maximized the silhouette score. This strategy revealed one group containing all wild-type, synonymous, and benign variants, two clusters of “DCM-like” variants (defined as Contractility Duration/Calcium Duration less than wild-type) and three “HCM-like” clusters (defined as Contractility Duration/Calcium Duration greater than wild-type) (Figure 4b,c). One of the DCM-like clusters only contained one variant, thus limiting the analysis for this group. Examining the contributions of each variable by cluster revealed that, while the calcium sensitivity metrics separated DCM-like and HCM-like clusters, there was also one HCM-like cluster, cluster 4, that had the highest values of calcium sensitivity (Figure 4d,e). Additionally, the HCM-like clusters were split by two variables relevant to diastole. The proportion of the maximum force that was contributed by the diastolic tension was increased in all HCM-like clusters, but cluster 1 showed less of an increase compared to clusters 4 and 5 (Figure 3f). Similarly, the relaxation rate for cluster 1 was slightly decreased, but clusters 4 and 5 had a more significant decrease in this rate (Figure 3g). Upstroke time of calcium did not distinguish between the HCM groups, but did separate the DCM-like groups, although this analysis was limited by the small number of DCM variants (Figure 3h). This data opens the possibility that there are multiple functional subclasses of HCM-like variants, demarcated by their relative calcium sensitivity and the degree of the associated diastolic dysfunction.

### Correlation of Clusters with Patient Outcomes

Given that the clusters we identified had clear functional differences *in vitro*, we asked whether the clusters correlated with clinical presentation. To answer this question, we collected a single-institution cohort of cardiomyopathy probands carrying pathogenic TNNI3 variants and asked whether our identified clusters correlated with clinical severity, as measured by an earlier age of onset. Our cohort only contained patients harboring HCM-like variants, and only two patients from the HCM-like-5 cluster, so we restricted our initial inter-cluster analysis to HCM-like-1 and HCM-like-4. We noted that patients carrying variants from HCM-like-4 presented significantly earlier than those carrying variants in HCM-like-1 (Figure 5a,b). To determine what was driving this difference, we then ran linear correlations with the five previously identified clustering metrics on the whole patient data set to determine which were significantly correlated with age of onset. Measures of diastolic function, most prominently the proportion of total force exerted in diastole, significantly correlated with age of onset (R^2^ of 0.6, Figure 5d, e). Interestingly, the previously examined measurements of calcium sensitivity, clear indicators of pathogenicity, did not significantly correlate with age of onset (Figure 5f,Figure S3). This raises the intriguing possibility that, in HCM, increased calcium sensitivity is necessary for cardiomyopathy to arise, but that the degree of resultant diastolic dysfunction determines the clinical severity of disease.

**Figure 5:**
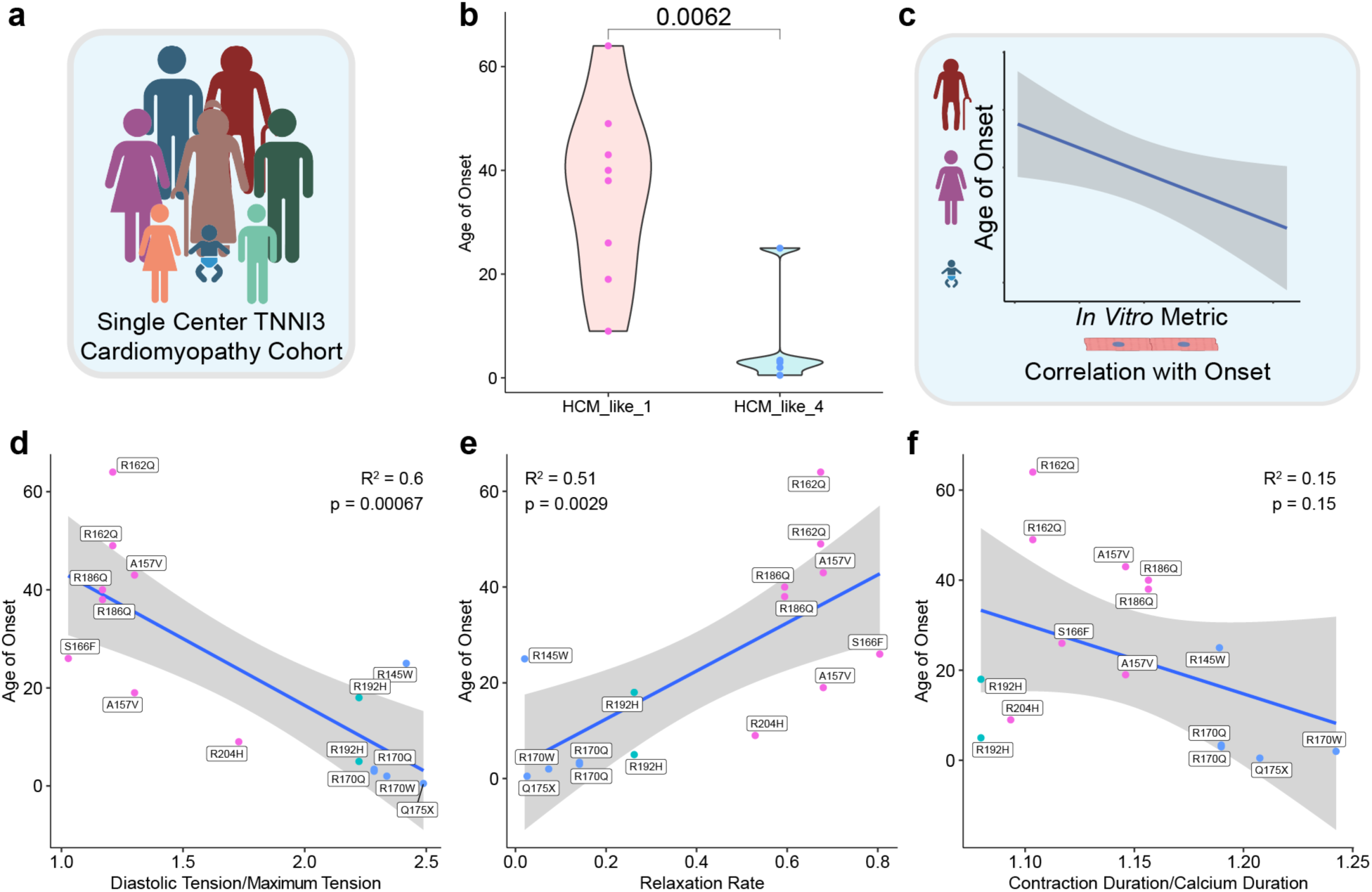
Correlation of Clusters to Clinical Severity. **a.** A single center cardiomyopathy cohort with TNNI3 variants was collected (N = 15 probands). **b.** Comparison of age of onset between patients with variants in HCM_like_1 and HCM_like_4 (only 2 patients carried variants in HCM_like_5 and so this was excluded from this analysis). **c.** Schematic of comparison of *in vitro* data to age of onset. **d.** Linear regression of age of onset vs Diastolic Tension/Maximum Tension. **e.** Similar comparison of Relaxation Rate and **f.** Contraction Duration/Calcium Duration. Significance in **b.** calculated using Wilcoxon rank-sum test.

## Discussion

The primary advance of this study is a scalable functional phenotyping platform that combines CRISPR-engineered Troponin I–null hiPSC-CMs as a chassis for the controlled re-expression of individual TNNI3 variants in a common background, simultaneous traction-based contractility and calcium imaging on substrata of physiological stiffness, and a computational pipeline that extracts more than 30 metrics and uses machine learning to select the most informative features for variant clustering. Leveraging this technical framework, we discerned HCM/RCM and DCM variants based on calcium sensitivity and force measurements, defined functional subgroups of disease variants, and showed that age of onset differed between HCM subgroups, suggesting that the functional metrics discerned *in vitro* may also impact clinical outcomes.

Our approach complements and extends previous studies that evaluate the pathogenicity of disease variants. Although multiple studies have analyzed sarcomeric cardiomyopathy variants using either purified proteins or stem-cell based models^15, 38–46^, the number of variants analyzed have been too few to provide more detail beyond binary assignment of pathogenicity/non-pathogenicity. More recent studies have expanded the number of variants analyzed by using gene expression or cell viability as a readout to suggest pathologic alterations leading to disease^13, 14^. However, since these studies only assessed a limited number of variables for each variant, they too were limited in their ability to evaluate clinical severity. Thus, the limited scale and parameter space of previous studies prevented an assessment of whether *in vitro* abnormalities correlate with disease severity. Our study addresses this critical gap by building a platform for multi-dimensional analysis of cardiac function applied to large numbers of variants.

Using our platform, we identified six clusters of variants, including 3 HCM-like clusters, that differed in their functional signatures. Interestingly, patients carrying variants from one cluster, HCM-like 1, had a later age of onset in our cohort compared to the other two. This appeared to be driven by the relatively lower degree of diastolic dysfunction in this group. This finding is consistent with clinical observations, as diastolic dysfunction is associated with significant morbidity and mortality in cardiomyopathy^47–52^. Interestingly, metrics measuring calcium sensitivity could reliably distinguish pathogenic from benign variants, but did not correlate with severity. This suggests a model where increased calcium sensitivity initiates disease, while other factors modify this initial insult to determine the ultimate diastolic function, which in turn determines the severity of the clinical presentation. Such factors could include incorporation of TNNI3 into the sarcomere, sarcomere organization, or protein stability, among others.

Several studies examine whether pharmacologic therapies that can alleviate pathology in models of HCM or RCM^12, 15, 46, 53^. However, these often examine a handful of models and do not directly correlate *in vitro* findings with disease severity. We found that two metrics, the ratio of diastolic tension to maximum tension and the relaxation rate, correlated with the age of onset in our cohort. These two metrics thus serve as excellent readouts in future pharmacologic studies attempting to identify therapies that ameliorate disease severity.

There are several limitations to this study. First, the platform quantifies cardiomyocyte-autonomous mechanisms (primarily contractile function and calcium cycling) and does not currently allow for the integration of other pathogenic mechanisms such as fibrosis that are thought to be important in cardiomyopathy^54, 55^. Further, the assay, as currently configured, evaluated single overexpressed variants while patients are predominantly heterozygous. Lack of heterozygosity might over-represent the impact of certain variants or miss potential interactions between alleles. This platform also does not capture other sources of genetic variation that may affect the exact clinical presentation of an individual patient, such as was recently seen with low-penetrance sarcomeric variants^56^. Finally, given the relative rarity of TNNI3-cardiomyopathy in the general population, our single-center cohort was limited in size. Due to this limitation, it lacked enough patients for thorough analysis of several of the identified groups, especially the DCM subgroups identified in our analysis.

In conclusion, we developed a scalable, multiparametric pipeline for analyzing the functional effect of *TNNI3* variants on cardiomyocyte function, and used it to not only distinguish between pathogenic and benign variants, but also to divide variants into subgroups with different clinical severity of disease. Additionally, we identified variables that linearly correlated with patient age of onset, suggesting that these measures may be useful in future studies identifying drugs that ameliorate disease presentation. This multidimensional approach will enhance our ability to sort phenotypically similar patients into functional and mechanistic categories, allowing for the development of personalized, genotype-informed treatments for cardiomyopathy.

## Non-Standard Abbreviations

hiPSC-CMs: Human Induced Pluripotent Stem Cell-derived Cardiomyocytes
DCM: Dilated Cardiomyopathy
HCM: Hypertrophic Cardiomyopathy
RCM: Restrictive Cardiomyopathy
DKO-CMs: TNNI1/TNNI3 Double Knockout Cardiomyocytes

## Acknowledgements

Figures 1,2,and 5 were created in part using BioRender.

## Supplementary Material

**Figure S1:**
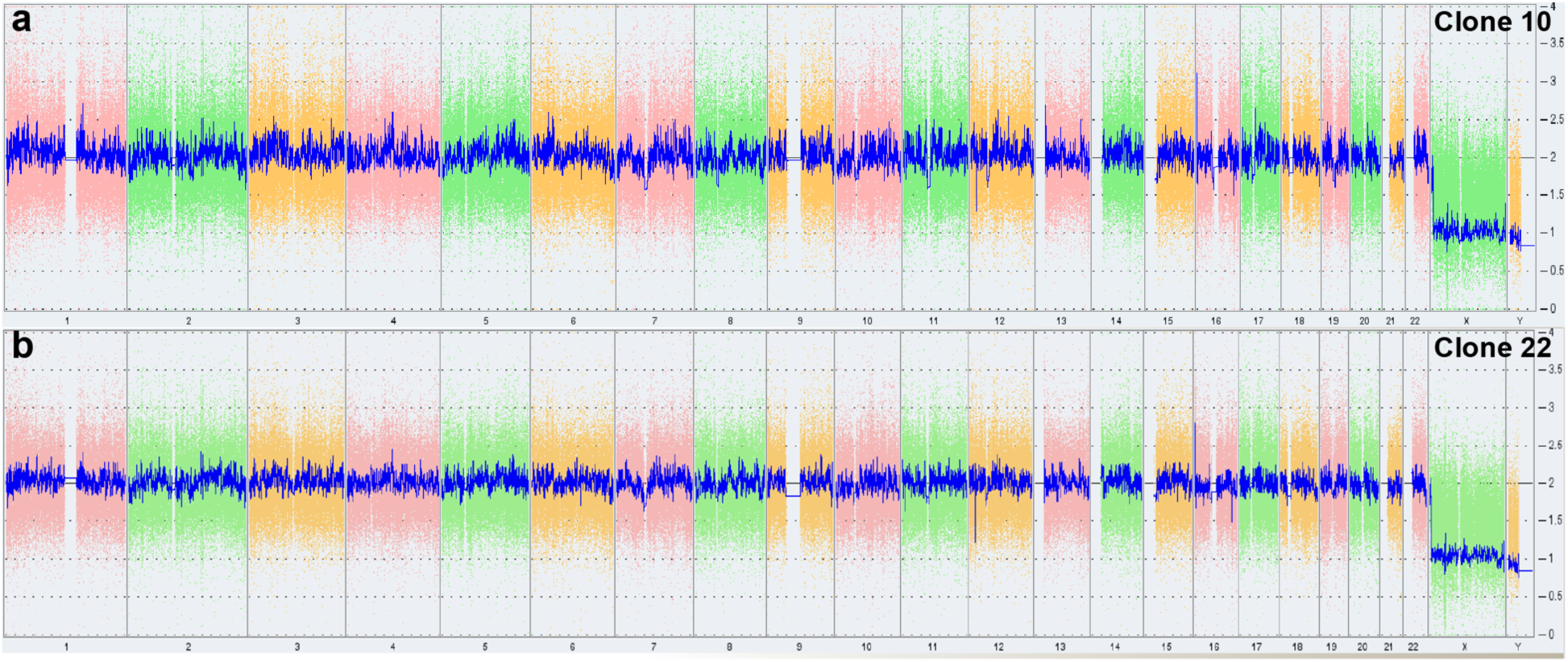
CNV Analysis of TNNI1/TNNI3 double knockout clones with Karyostat. **a.** Clone 10 and b. Clone 22 show normal Karyostat results.

**Figure S2:**
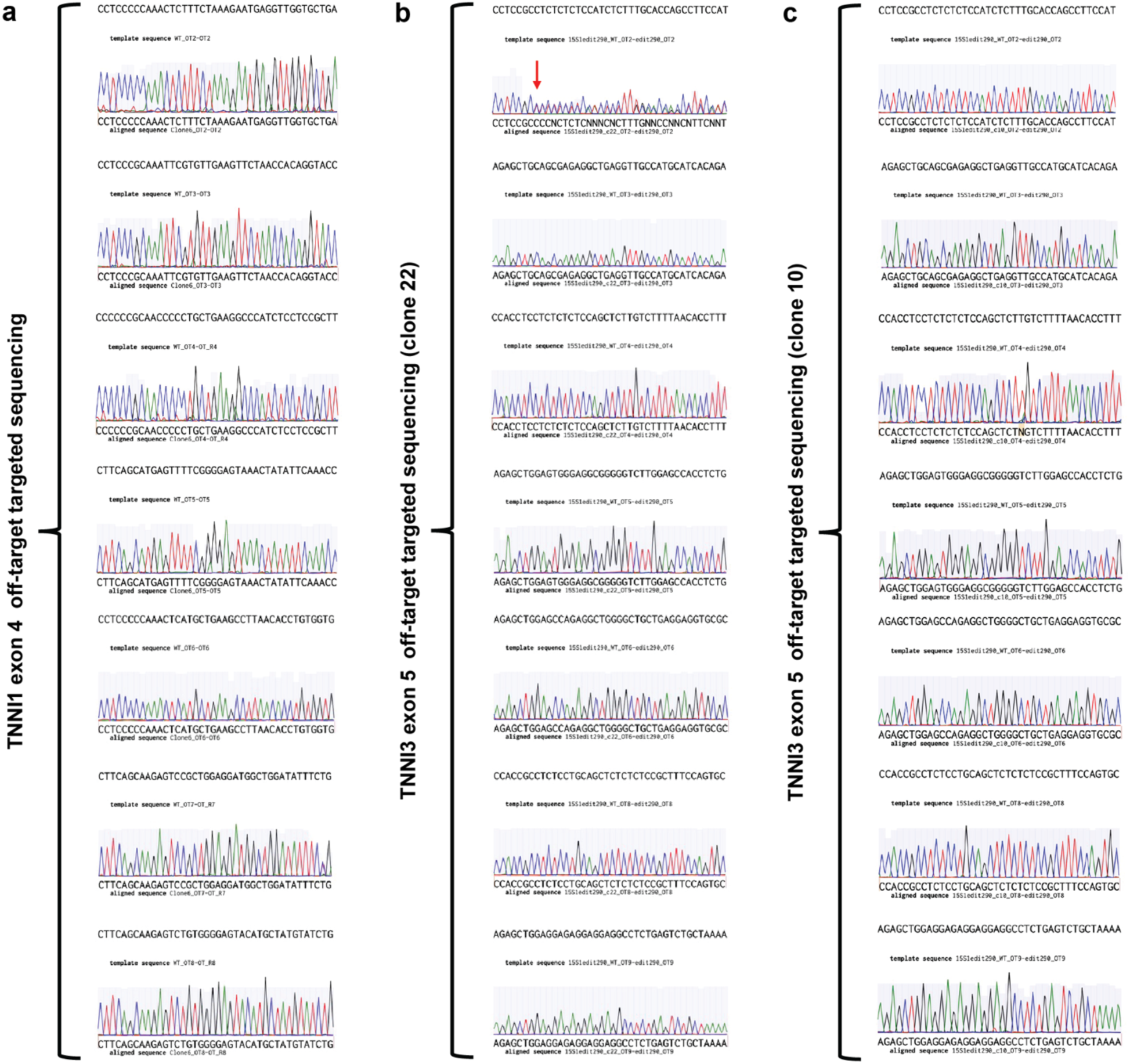
Off-target analysis of TNNI1/TNNI3 double knockout clones. **a.** Off target sequencing of parental TNNI1 locus in TNNI1 KO line. WT sequence shown at the top of each trace. Trace represents sanger sequencing of each off-target locus identified by COSMID ^23^. **b.** Off target sequencing of the TNNI3 locus in clone 22. Heterozygous frame shift denoted with red arrow. c. Off target sequencing of TNNI3 in clone 10 shows no off-target edits.

**Figure S3:**
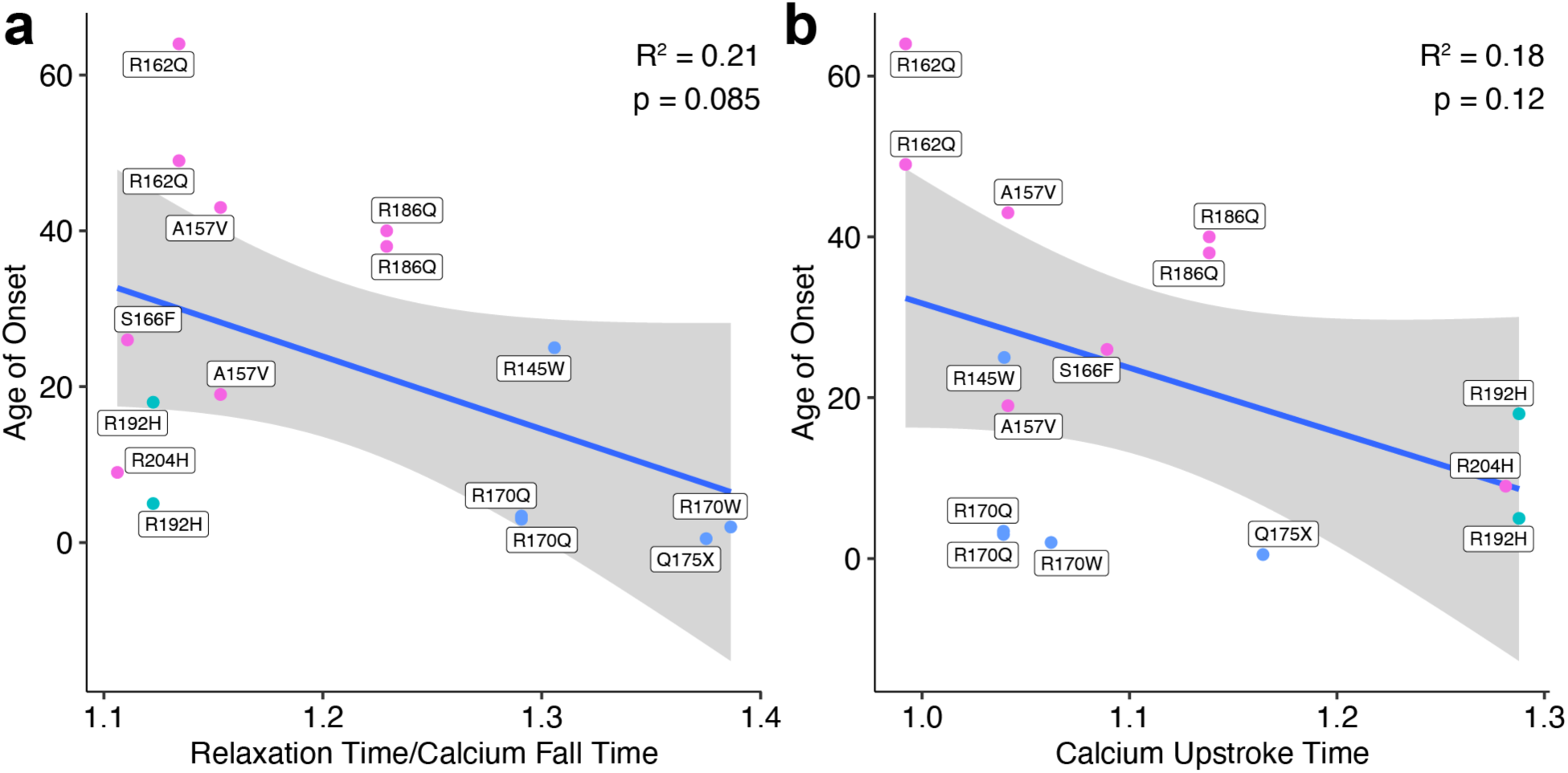
Correlation with Additional Clustering Variables. **a.** Linear regression of age of onset vs Relaxation Time/Calcium Fall Time and **b.** vs Calcium Upstroke Time

